# The Pleiotropy Hypothesis of Molecular Evolution

**DOI:** 10.1101/2025.11.19.689358

**Authors:** Xun Gu

## Abstract

Under the nearly-neutral model of protein evolution, the evolutionary rate is virtually determined by the selection intensity (*S*), which can be decomposed into *S*=-*K*×*B_0_*, where *K* is the gene pleiotropy defined by the number of fitness-related traits (molecular phenotypes) and *B_0_* is the baseline of the selection intensity. Hence, the variation of *S* (sequence conservation) among genes may have two resources: one is the variation of gene pleiotropy among genes (*K*-mode), and the other is the variation of baseline intensity among genes (*B*-mode). While *K* can be effectively estimated (denoted by *K_e_*) based on the phylogenetic analysis of protein sequences, the correlation between *K_e_* and empirical pleiotropy measures remains uninvestigated. In this paper, we show positive correlations of effective gene pleiotropy with protein-protein interactions, expression broadness, enzyme connections, and involved biological processes. We thus propose the pleiotropy hypothesis (*K*-mode), suggesting that the rate variation among proteins is mainly due to the variation of gene pleiotropy, revealing a sophisticated display of multiple gene functionality.

## Introduction

PLEIOTROPY, or the capability of a gene to affect multiple to affect multiple traits plays a central role in genetics, development, and evolution (Fisher 1930; Wagner and Zhang 2011; Paaby and Rockman 2013). Like many biological terminologies, the concept of pleiotropy is easy to understand but difficult to measure. Practically, biologists address this issue by experimentally identifying distinct phenotypes, in vitro or in vivo, caused by the same mutation (mutational pleiotropy) or mutations from the same gene (gene pleiotropy). Although these studies largely focused on the nature of phenotypes, the results did provide a “minimum” estimate about the extent of pleiotropy. Thus, one may compare whether a gene is more pleiotropic than others, as long as at the same technological platform.

However, different pleiotropy measures based on different technological platforms certainly represent biologically significant differences that may even lead to contradictory interpretations. Paaby and Rockman (2013) formulated three types of gene pleiotropy. (*i*) Molecular pleiotropy (*f*) refers to the number of protein functions most likely in vitro. (*ii*) Developmental pleiotropy (*d*) provides concrete examples to demonstrate how a single gene or mutation may affect distinct phenotypes *in vivo*. And (*iii*) selectional pleiotropy (*n*) refers to the number of molecular phenotypes (Gu 2007), each of which corresponds to a single nontrivial fitness component; in this sense, it is also called fitness-related traits. While the difference between *f* and *d* illustrates the many-face problem of pleiotropy, the difference between *n* and *f* (or *d*) represents the conceptual gap between the empirical pleiotropy and fitness.

Although functional genomics has brought high-throughput data to bear on the nature and extent of pleiotropy (Dudley et al. 2005; Ohya et al. 2005; Pal et al. 2006; Cooper et al. 2007), this issue remains highly controversial, largely because of the phenotypic complexity. In this article, we address this issue by investigating the relationship between empirical pleiotropy and the selection (fitness-related) pleiotropy. While the former is based on the measurement of affected phenotypes, we suggest the use of the later should be based on the estimation of protein sequences to remove the artifact caused by data correlations. Gu (2007) developed a statistical method to estimate the “effective pleiotropy” (*K_e_*) of a gene from the multiple-sequence alignment (MSA) of protein sequences. Most genes have *K_e_* in the range between 1 and 20 (Su et al. 2010; Zeng et al 2010), with the medium *K_e_*=6.5 of these estimates that is comparable to some empirical pleiotropy measures (Wagner and Zhang 2011; Paaby and Rockman 2013). Further, Gu (2014) showed that *K_e_* has an intrinsic relationship with the rank of the genotype–phenotype map, and derived the condition that *K_e_* can be interpreted as effective gene pleiotropy. The goal of this paper is to establish the correlation between *K_e_* and several empirical pleiotropy measures. Our hypothesis is schematically summarized in **Fig.1**.

**Fig. 1.**
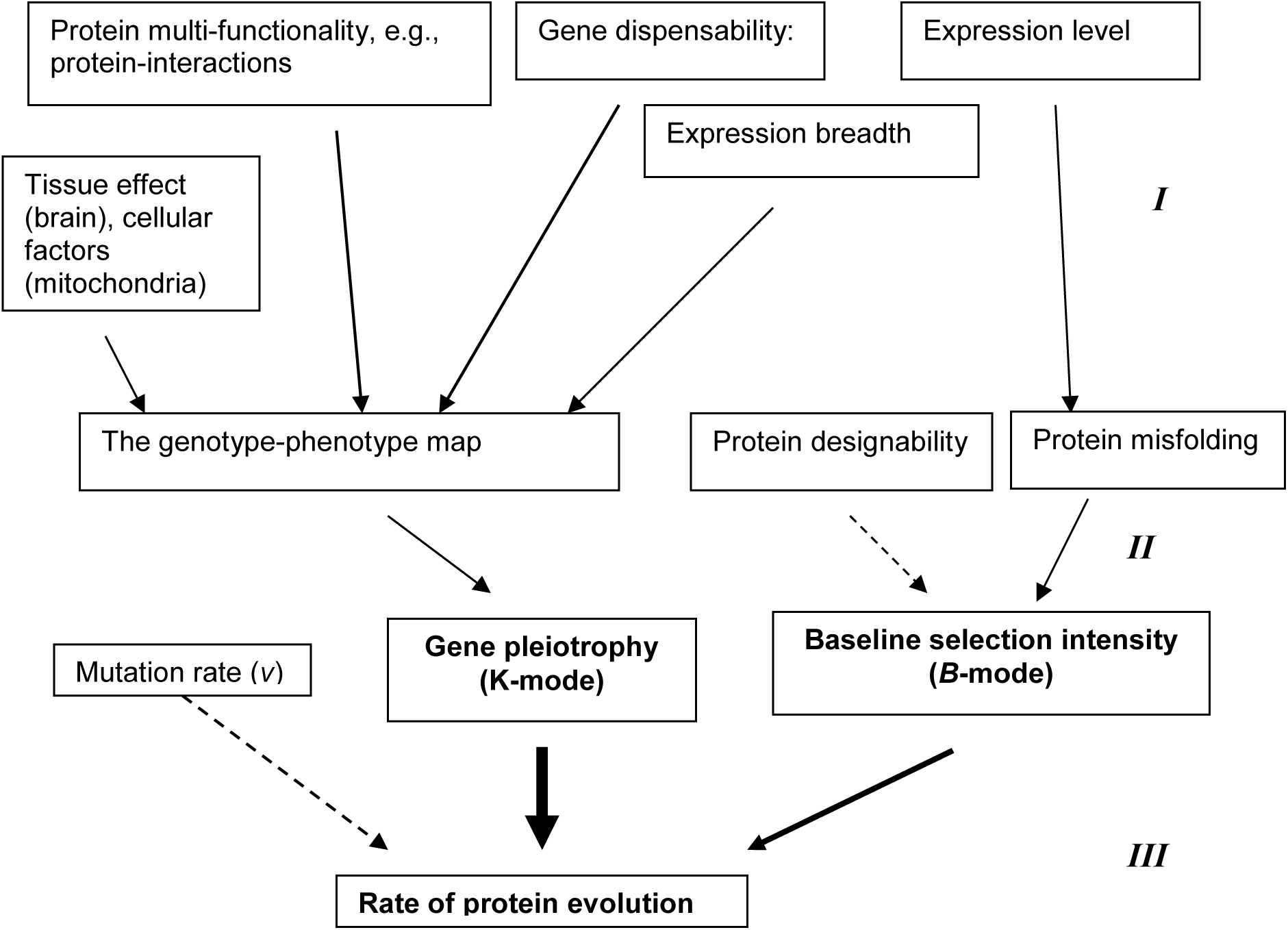
A schematic presentation of the K-B hypothesis about the genomic correlations with protein evolution. Up-level system factors are grouped into two categories: Roles in genotype-phenotype mapping, and molecular properties (Level-I), corresponding to *K* and *B*-modes in the pleiotropy model of protein evolution (Level-II). At level-III, together with the mutation rate (dashed-line), these two modes determine the rate of protein evolution.

## Results

### The pleiotropy hypothesis of molecular evolution: a theoretical argument

The theory of molecular evolution postulates that the evolutionary rate (λ) of a nucleotide site is determined by three layers of genetic processes: mutation at the individual level, polymorphism at the population level, and fixation at the species level (Kimura 1983). Let *v* be the mutation rate, *s* the coefficient of selection and *N_e_* the effective population size. It has been shown that the evolutionary rate is given by

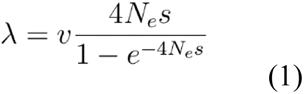

Eq.(1) provided a solid foundation to distinguish between adaptive evolution, neutral evolution, and nearly-neutral evolution: it predicts λ/*v*>1 for adaptive evolution (*s*>0), λ/*v*=1 for neutral evolution (*s*=0), or λ/*v*<1 for deleterious evolution (*s*<0), respectively.

Let *S=4N_e_s* be the selection intensity that uniquely determines the substitution-mutation rate ratio λ/*v.* Eq.(1) indicates that variation of substitution rates among genes is determined by the variation of selection intensity (*S*) among genes. Under the theme of nearly-neutrality without adaptive selection, the view of genotype-phenotype mapping (Gu 2007; 2014) demonstrated that the structure of *S* can be written as the product of two components, that is

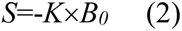

where *K* is the rank of genoty pe-phenotype mapping as a proxy of gene pleiotropy and *B_0_* is the baseline selection intensity; the negative sign indicates the condition of *S<*0 (near-neutrality). One may therefore envisage (Zeng et al. 2009) two selection modes: the variation of *S* among genes (or the sequence conservation) is attributed to the variation of gene pleiotropy among genes (the *K*-mode), or to the variation of baseline intensity among genes (*B*-mode). The pleiotropy hypothesis of molecular evolution claims that the *K*-mode is more dominant than the *B*-mode, that is, the substitution rate is mainly determined by the degree of gene pleiotropy, whereas the baseline selection intensity remains a rough constant.

### A large variation of effective gene pleiotropy among genes

Gu (2007) developed a statistical method to estimate the effective gene pleiotropy (*K_e_*) and the baseline selection intensity *B_0_* effectively, based upon the phylogenetic analysis of protein sequences. One may see Data and Methods for a brief introduction: *K_e_* and *B_0_* can be estimated from *d_N_/d_S_* (the ratio of nonsynonymous to synonymous rates) and *H* (a relative measure for rate variation among amino acid sites). Based on limited datasets, our previous studies () have revealed a large variation of *K_e_* among genes and a small variation of *B_0_*. Here we confirm this conclusion by two large datasets: yeasts and vertebrates.

**Fig.2A** shows the *d_N_/d_S_*∼(1-*H*) plotting of 3470 yeast genes, where almost all genes are below the line of *K_e_*=1, suggesting that *K_e_*>1 holds virtually for those genes. The histogram of *K_e_* of yeast genes (**Fig.2B**) reveals that most genes have substantial degrees of pleiotropy (*K_e_*>2), and the mean of effective gene pleiotropy is *K_e_*=7.76 and the 25%-75% quantile is from 4.2 to 8.8. Besides, we estimated the baseline selection intensity *B*_0_ for those yeast genes. In contrast to the broad variation of effective gene pleiotropy, the baseline selection intensity only has a mild variation among genes, with the mean around 1.3, with the 25% to 75% quantile (0.96, 1.78). Overall, the variation of *K_e_* among yeast genes may explain up to 75% variation of the nonsynonymous distances among genes (**Fig.1C**).

**Fig. 2.**
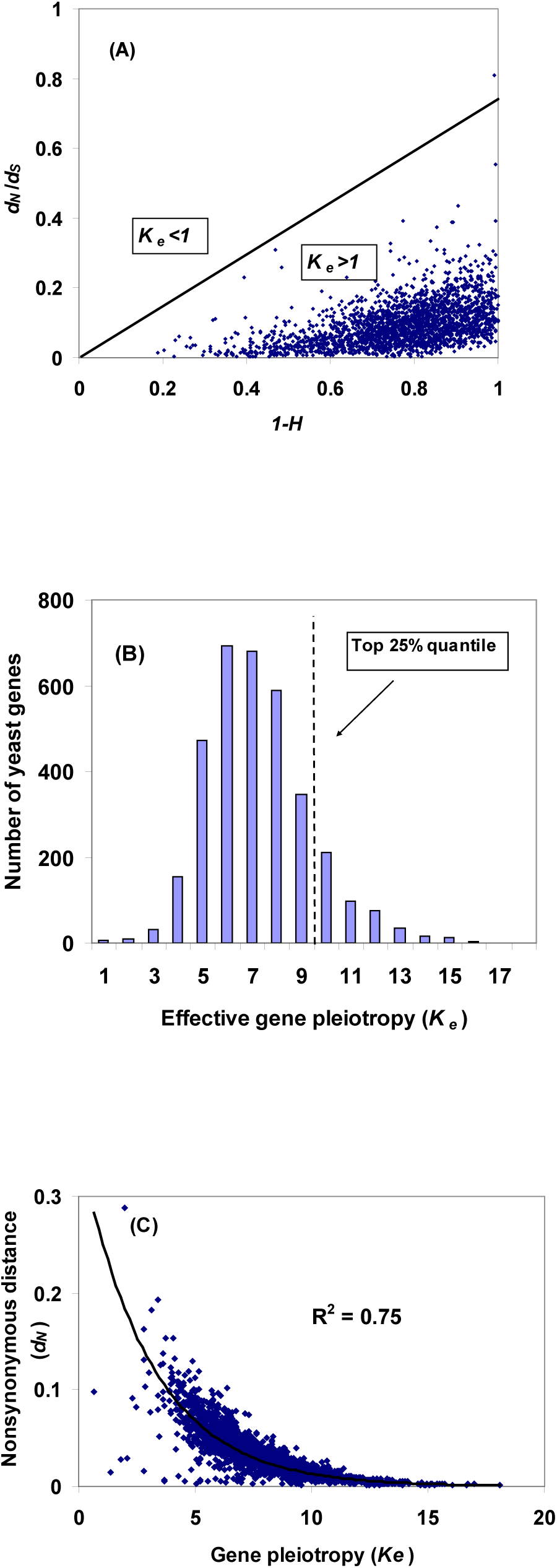
Extent of gene pleiotropy and baseline selection intensity. (**A**) The *d_N_/d_S_* ∼ (1-*H*) plotting for 3470 yeast genes. The straight lines indicates the areas of *K_e_* <1 (biologically unrealistic) and *K_e_* >1, respectively. For each gene, the *d_N_/d_S_* ratio was estimated by the orthologs of *S. cerevisiae* and *S.bayanus*, while the index *H* was estimated from five yeasts. (**B**) The histogram of effective gene pleiotropy (*K_e_*) for 3470 yeast genes. (**C**) The nonsynonymous distance (*d_N_*) plotting against the effective gene pleiotropy for 3470 yeast genes. *Data resources*: in total 3470 yeast genes were analyzed, based on five yeasts (*S. cerevisiae*, *C.glabrata*, *K.lactis*, *D.hansenii* and *Y.lipolytica*). For each gene, the *d_N_/d_S_* ratio was estimated by two closely-related yeasts, *S. cerevisiae* and *S. paradoxus,* and the *H-*index based on the yeast phylogeny of five species.

Similarly, we analyzed 4336 vertebrate genes. For each gene, the *d_N_/d_S_* ratio was obtained from the human and mouse, while the *H*-index was estimated by a given vertebrate phylogeny that includes eight vertebrate species (Gu 2007). We found that the mean of *H* among vertebrate genes is 0.517. Most of those genes show a certain degree of pleiotropy (*K_e_*>1), supporting the notion that gene pleiotropy is a general pattern. The mean of *K_e_* (6.52) indicates that a vertebrate gene typically affects 6∼7 molecular phenotypes (Gu 2007; Gu 2014).

### Weak correlation of *K_e_* with protein-protein interactions

The pleiotropy hypothesis of molecular evolution predicts that the effective gene pleiotropy *K_e_* would have a positive correlation with the degree of protein-protein interactions, under the premise that proteins with high interactivity tend to be involved in many biological processes, i.e., high gene pleiotropy. Three yeast protein-protein interaction datasets were used to test this prediction; see the legend of Fig.3 for data description, as well as the empirical criteria of highly pleiotropic genes. The main prediction we try to test is that highly pleiotropic genes should appear more frequently in hubs than single-link genes or nonhubs. To this end, we calculated *F_P_* for each interaction group, the fraction of highly pleiotropic genes, by the proportion of genes that have *K_e_*≥9. **Fig.3A** shows that in three protein interaction datasets, *F_P_* in the hub group is significantly higher than that of non-hubs (1.6-fold) or singlish-link genes (2.0-fold), *P-*value<10^-6^, suggesting that a hub gene tends to be highly pleiotropic.

**Fig. 3.**
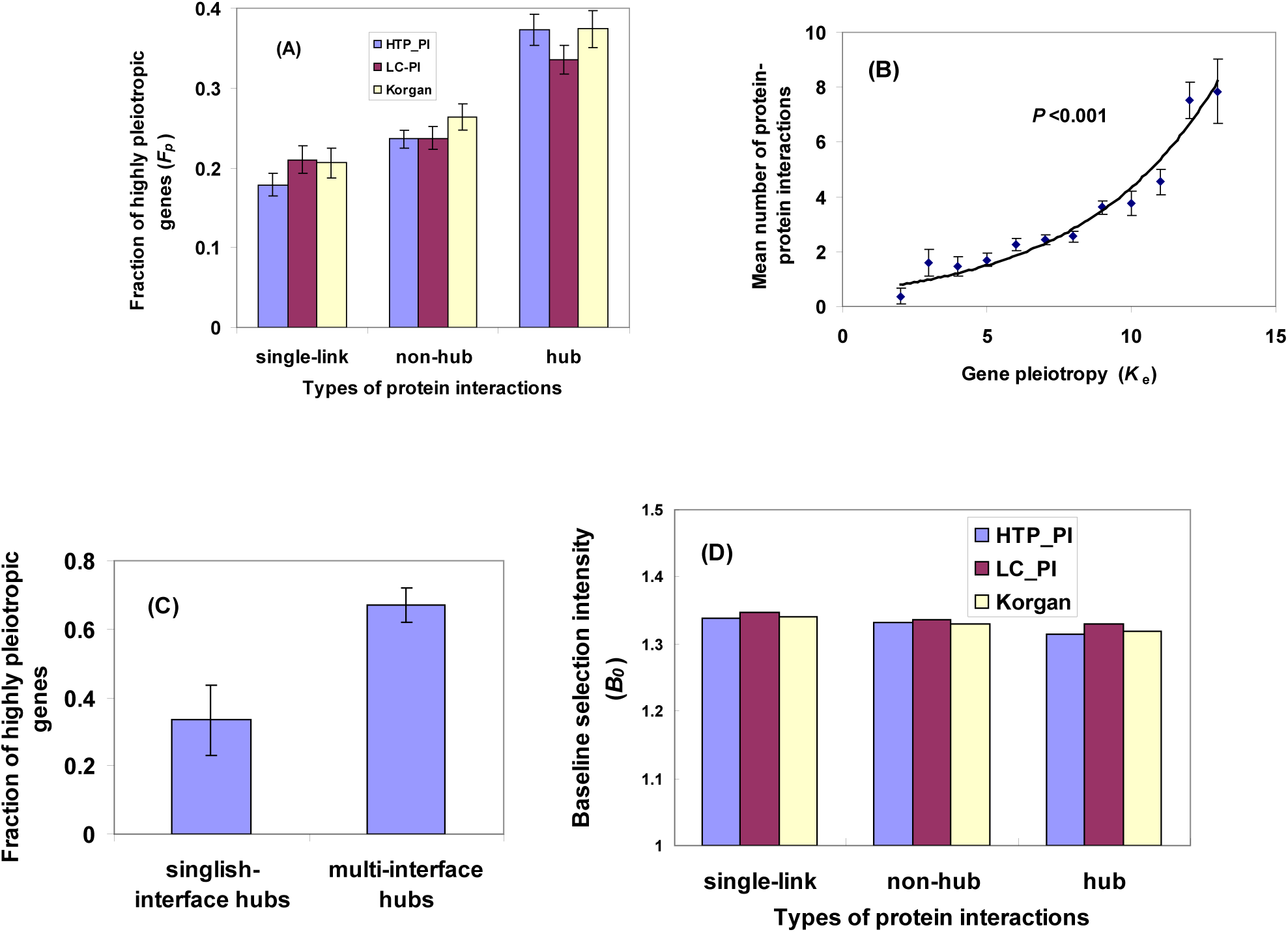
Gene pleiotropy (K-mode) and protein-protein interactions. (**A**) Fraction of highly pleiotropic genes (*F_P_*), defined as the proportion of yeast genes with *K_e_*≥9 (25% top), is significantly higher in hub genes than that in non-hub genes or single-link genes. Three years protein interaction datasets used: HTP_PI (high throughput, 2677 genes), LC_PI (literature curated, 2122 genes), and Korgan (1621 genes). (**B**) Correlation between number of interactions (HTP_PI) and the degree of pleiotropy. (**C**) Yeast protein interface-interaction network: *F_P_* is higher in multi-interface-hubs than that in singlish-interface-hubs; the classification of interface-hubs from the original paper. (D) The mean of baseline intensity (*B_0_*) is invariant among hub types as well as data types. *Criteria*: define a highly pleiotropic gene by the cutoff *K_e_*≥9, roughly corresponding to the top 25% quantile (see Fig.2B) of the pleiotropic histogram among yeast genes. For each protein-protein interaction dataset, genes were classified into single-link genes, nonhubs (<7 links) or hubs (≥7 links), respectively. Note that variations in cutoffs for both pleiotropy and connectivity did not affect our results.

In another analysis, we grouped genes with similar gene pleiotropy, and calculated the mean number of protein-protein interactions and standard errors. **Fig.3B** shows the result based on the HTP-PI dataset, indicating that the mean number of interactions increases with the degree of gene pleiotropy; *P-value*<0.001. This so-called pleiotropy-connectivity correlation was further tested by the structural interface dataset of protein-protein interactions (Kim et al 2006). We followed the treatment used by the original authors to classify yeast hub genes into two groups: singlish-interface hubs (one or two structural interfaces) and multi-interface hubs (at least three interfaces). **Fig.3C** showed a higher *F_P_* in hubs with multiple interfaces than hubs with singlish interface (*P-value*<0.01). By contrast, we found that the mean of baseline intensity (*B_0_*) is almost invariant among hub types as well as data types (**Fig.3D**). A previous study (Kim et al. 2006) pointed out that those protein interaction datasets led to inconsistent conclusions about the correlation between the rate of protein evolution and the protein connectivity.

### Relationship between gene pleiotropy, tissue broadness and tissue specificity

In multiple-tissue organisms, between the organismal fitness and genes is the hierarchy of biological systems that may shape the evolutionary rate and the degree of gene pleiotropy simultaneously. Functioning of a protein in diverse tissues (*expression breadth*) is one of major mechanisms for gene pleiotropy, implying that gene pleiotropy may underlie the rate-expression breadth correlation (Pal et al. 2006; Duret and Mouchiroud 2002; Su et al. 2004). We used the human RNA-seq data to test this notion. A gene is classified into the *H*-group (high) of tissue broadness if the number of expressed tissues is over 75% quantile), or otherwise into the *L*-group (low). As shown by Table 1, the fraction of highly pleiotropic genes *F_P_* of the *H-*group is significantly higher than that in the *L*-group (χ^2^-test, *P*<10^-5^), while the baseline selection intensity *B*_0_ is virtually the same between the *H* and *L*-groups. We thus conclude that human genes expressed in more tissues tend to be more pleiotropic.

**Table 1.** Analysis of tissue-specificity, broadness and gene pleiotropy.

| Types | Brain-specific | Not-brain specific | Not-specific |
| --- | --- | --- | --- |
| Gene number | 331 | 1980 | 12063 |
| Expression broadness | $32.7 \pm 1.3$ | $31.8 \pm 0.6$ | $42.2 \pm 0.3$ |
| Gene number | 73 | 390 | 3186 |
| Gene pleiotropy | $8.18 \pm 0.39$ | $6.14 \pm 0.13$ | $6.61 \pm 0.05$ |
| $F_p$ | $0.55 \pm 0.06$ | $0.19 \pm 0.02$ | $0.25 \pm 0.01$ |

**Table 2.** Gene pleiotropy analysis of mitochondrial proteins.

| | Num of genes | Effective pleiotropy ($K_e$) | | | Frac. highly pleiotropic genes ($F_p$) |
| --- | --- | --- | --- | --- | --- |
| | | Mean ± s.e. | Medium | Quantile (25%-75%) | |
| Non-mit genes | 3993 | 6.64±0.04 | 6.05 | 4.51-7.93 | 24.3±0.7 % |
| Mit genes (Swiss) | 131 | 5.16±0.17 | 5.24 | 3.69-6.56 | 8.3±2.4 % |
| Mit genes (mitop2) | 337 | 5.19±0.12 | 4.85 | 3.69-6.32 | 9.2±1.6 % |
| mtEnzyme | 179 | 5.15±0.15 | 4.91 | 3.81-6.31 | 8.4±2.1 % |
| mtDNA-proc | 57 | 4.37±0.27 | 4.07 | 3.26-5.10 | 1.8±1.7 % |
| mtAssembling | 83 | 5.91±0.25 | 5.74 | 4.40-7.18 | 16.9±4.1 % |
| Ribosomal proteins |  |  |  |  |  |
| Mitochondrial | 19 | 4.68±0.48 | 4.26 | 3.45-6.30 | 5.3±5.2 % |
| Cytosolic | 20 | 11.1±1.20 | 9.96 | 6.38-16.5 | 65±11 % |

On the other hand, the fitness effect of a single tissue could be substantially multiplied such as tissue brain in vertebrates. We addressed this issue below. Technically, a gene is called brain-specific if it has the highest expression level in brain, which is at least two-fold higher than that in other tissues. With the same criterion, a gene is called not-brain tissue specific if the tissue with the highest expression level is rather than brain. Finally, a gene is called broadly-expressed if it is not tissue-specific in any tissue we examined. For 3649 human genes we examined, we found, strikingly, that the fraction of highly pleiotroic genes in (not-brain) tissue-specific genes (*F_P_*=19.2%) is significantly lower than that (*F_P_*=25.0%) in broadly-expressed genes (*P-value<*0.05), whereas *F_P_* in the brain-specific genes is extraordinarily higher (54.8%); *P-value*<0.001. Our finding suggests a strong relationship between the brain function and the fitness of organism may shape the pleiotropy of genes that function mainly in the brain (Table 1).

### Correlation of *K_e_* with gene essentiality

The relationship between the rate of protein evolution and gene essentiality, short for the rate-essentiality correlation (Wilson et al. 1977), remains controversial for the underlying mechanism (Pal et al. 2003; Krylov et al. 2003; Wall et al. 2005). We propose that gene pleiotropy underlies the rate-essentiality correlation, claiming (*i*) that a (single-copy) dispensable gene implies effective genetic buffering (Gu 2003), that (*ii*) the likelihood for being dispensable is low for a highly pleiotropic gene because all affected fitness components have to be genetically buffered, and that (*iii*) highly pleiotropic genes evolve slowly. Indeed, we observed that the fraction of highly pleiotropic yeast genes in the indispensable (essential) group (*F_P_*≍35%) is about 1.5-fold higher than that in the dispensable group (*F_P_*≍23%, *P-value*<10^-8^), whereas the mean of baseline selection intensity is almost the same (*B*_0_=1.34 and 1.33, respectively).

Further, 1768 yeast single-copy genes were grouped according to *K_e_*=1, 2,…, and calculated *P_E_,* the proportion of essential genes, in each group. **Fig.4A** shows a significant correlation between dispensability (*Q*=1- *P_E_*) and gene pleiotropy (*K_e_*) (*r*=-0.90, *P*<0.001), whereas *Q* is independent of *B*_0_ (**Fig.4*B***). Clearly, a highly pleiotropic gene tends to have a high chance to be essential. This pattern is compatible with a simple model: suppose that a (single-copy) gene is dispensable when all *K* molecular phenotypes of the gene is genetically buffered when this gene is deleted. Let *q* be the fidelity of genetic buffering for a single molecular phenotype. Hence, the proportion of dispensable gene (*Q*) is given by *Q=*1- *P_E_=q^K^*, which fits the yeast single-gene deletion fitness data well (**Fig.4A**). Interestingly, we estimated *q*≈0.94, indicating that the fidelity of genetic buffering for a single molecular phenotype (fitness-related trait) is fairly high, or the error rate is small. However, for highly pleiotropic genes, error accumulation in genetic buffering could become nontrivial, increasing the change to be essential, which can explain the inverse rate-essentiality correlation.

**Fig. 4.**
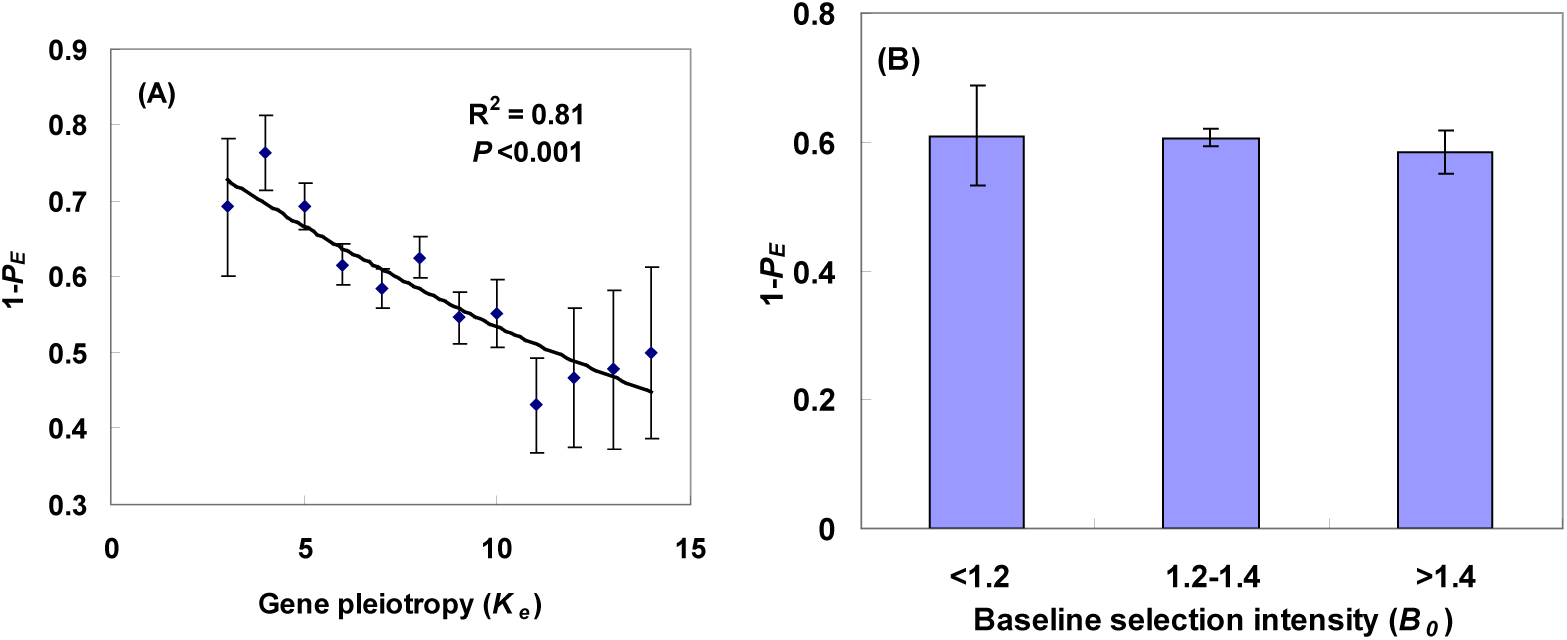
The Pleiotropy hypothesis of genetic robustness: (A) *Q*=1- *P_E_* plotting against the gene pleiotropy (*K_e_*), where *P_E_* is the proportion of essential genes. (B) *Q*=1- *P_E_* plotting against the baseline selection intensity (*B_0_*).

### Mitochondrial proteins have low pleiotropy due to functional modularity

Gene pleiotropy could be constrained by the prevalence of modularity in multicellular organisms. However, to what extent the effect of modularity on pleiotropy remains an unsolved issue (Welch and Waxman 2003; Wagner et al. 2008). We used mammalian mitochondria as a model of modularity to study the effect of modularity on gene pleiotropy. In mammals, mitochondria are composed of hundreds of proteins (mitochondrial proteome), which are nuclear-encoded except for 13 mitochondrial-encoded proteins. One may anticipate that functionality of mitochondria may strongly limit highly pleiotropic genes in the mitochondrial proteome.

We compared the histograms of effective gene pleiotropy (*K_e_*) for 3993 non-mitochondrial proteins (genome genes) and 468 (predicted) mitochondrial proteins. The Kolmogrov-Smirnov (KS) test showed that they differ highly significantly (*P-value*<10^-6^). Roughly speaking, while a typical non-mitochondrial gene may affect 5-8 fitness components, a typical mitochondrial gene may affect 4-6 fitness components, according to the 25%-75% quantile of effective gene pleiotropy. We examined the pleiotropy of mitochondrial ribosomal proteins, as they evolve much faster than their cytosolic counterparts (Fig.5): the mean of effective pleiotropy of mitochondrial ribosomal proteins *K_e_*=4.68±0.48 is considerably lower than that of cytosolic ones (*K_e_*=11.1±1.2); *P-value*<10^-4^. As these two types of ribosomal proteins have virtually the same sequence-structure properties, we provide a concrete example to show that gene pleiotropy instead of the structural property determines the rate of protein evolution.

**Fig. 5.**
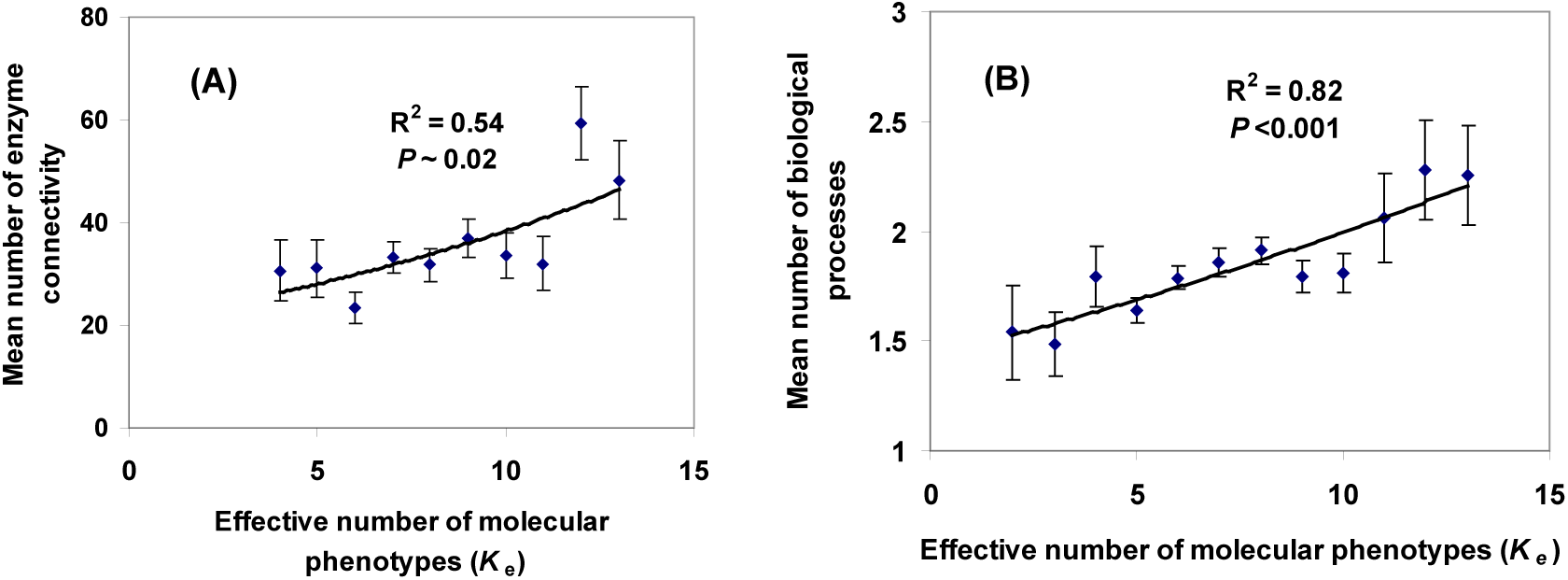
(A) A significant positive correlation between enzyme connectivity and the effective gene pleiotropy (*P*∼0.02). (B) a positive correlation between the number of biological processes and the mean of *K_e_* (*P*<0.001).

Moreover, we argue that the reduced pleiotropy in mitochondrial genes is mainly caused by the lack of highly pleiotropic genes, rather than the enrichment of low-pleiotropic genes. We then used *F_P_*, the fraction of highly pleiotropic genes defined by *K_e_*≥8 (roughly corresponding to the top 25% quantile), to catch this feature. As expected, in the non-mitochondrial gene set, *F_P_* corresponds to the top 24.3% quantile. Strikingly, we obtained *F_P_*= 9.20% (mitop2) or 8.3% (Swissprot) for mitochondrial proteins (Table 1), which is much lower than that of non-mitochondrial proteins (*P-value*<10^-6^, a 2×2 test). Our findings suggest that the effect of modularity is strong for highly pleiotropic genes.

One may wonder whether under-presentation of highly pleiotropic mitochondrial genes may depend on the functional preference of genes. To this end we tentatively classified mitochondrial proteins into three groups: (*i*) proteins and enzymes involved in the oxidative phosphorylation pathway of energy conversion and other metabolic processes (mtEnzyme group, 179 genes); (*ii*) proteins involved in mitochondrial DNA replication, transcription and translation (mtDNA-processing group, 57 genes) and (*iii*) proteins involved in signaling, transportation and membrane proteins for mitochondrial assembling and functioning (mtAssembling group, 83 genes). Intuitively, the pleiotropy of mtEnzyme genes may typically represent the multi-functionality of mitochondria. We further speculate that the pleiotropy of mtAssembling genes is ‘universal’ by the means of mitochondrial functions. In addition, mtAssembling genes tend to have multiple sub-cellular roles. By contrast, the pleiotropy of mtDNA processing genes should be considerably reduced because all these genes are only for 13 mitochondrial-encoded proteins plus several tRNAs and rRNAs. Together, we predict that the degree of pleiotropy should follow the order of mtDNA-processing < mtEnzyme < mtAssembling.

Strikingly, we have observed that the pleiotropy precisely follows the order predicted above (Table 1): The mean effective pleiotropy *K_e_* is 4.57 (mtDNA-processing), 5.15 (mtEnzyme) or 5.91 (mtAssembling); their differences are statistically significant (ANOVA, *P*<0.001). Similarly, the fraction of highly pleiotropic genes *F_p_* is 1.8% (mtDNA-processing), 8.4% (mtEnzyme) or 16.9% (mtAssembling); 2×3 table test, *P*<0.01. By gene ontology (GO), we compiled a list of genes involved in nuclear DNA processing (replication, transcription and translation) that has *K_e_* =6.84 on average, and *F_p_* is 26.9%. Impressively, small mitochondrial genome has prevented genes for mtDNA-processing from being highly pleiotropic. In particular here we report a clear-cut case (Table 1): Reflecting their universal roles in the organismal fitness, cytosolic (*cy*) ribosomal proteins are usually highly pleiotropic; most of them are on the top of 10% of all genes studied. By contrast, mitochondrial (*mt*) ribosomal proteins have low degrees of pleiotropy, around the bottom 25%. The average difference of effective pleiotropy between *cy* and *mt* ribosomal proteins is Δ*K_e_*=11.1-4.68=6.42 (Welcox test, *P*<0.001). Comparing to cytosolic ribosomal proteins, it is impressive that the fraction (*F_p_*) of highly pleiotropic genes in *mt* ribosomal proteins has been reduced more than one magnitude (5.5% *mt* versus 65% *cy*, *P*<0.001).

## Discussion

### The pleiotropy-expression correlation

It is well-known that the evolutionary rate and the expression level is strongly inversely correlated (Yang et al. 2003). The controversy is how to explain this pattern. There are two possibilities. (*i*) Expression level is the determinant of the rate of protein evolution (Drummond et al. 2005; 2006). And (*ii*) protein sequence conservation and expression level are both determined by the gene pleiotropy. In single-cell organisms, a highly pleiotropic gene tends to be highly expressed because it requires a large amount of protein molecules. We used two expression measures to test this prediction: yeast CAI (codon adaptation index) and yeast Abd (proteome-wide protein abundance) (Batada et al. 2006). We found a statistically significant positive correlation between *K*_e_ and CAI (*r^2^*=0.05, *P*<10^-6^), as well as between *K*_e_ and Abd (*r^2^*=0.07, *P*<10^-8^).

However, this issue could be complicated by the high correlation between the evolutionary rate and the expression level (Pal et al. 2006; Duret and Mouchiroud 2002). To address this issue, we used the synonymous distance (*d_S_*) between the human and mouse. Since a highly expressed gene tends to have a low *d_S_*, one can predict a negative correlation between *B*_0_ and *d_S_*, which is indeed the case (*R*=-0.12, *P*<10^-8^). Noting that *K* and *d_S_* has a very weak correlation (*R*=-0.02), we concluded that the effect of expression breadth on the rate of protein evolution is *K*-mode dependent, while the effect of expression level is *B*-mode dependent.

### Correlation of *K_e_* with biological processes

In spite of limited datasets, we analyzed the correlation of effective gene pleiotropy (*K_e_*) with the enzyme connectivity, as well as the number of biological processes involved in GO (gene ontology). For 318 yeast enzymes, we found a significant positive correlation between enzyme connectivity and the effective gene pleiotropy (*P*∼0.02) (**Fig.5, A**). For instance, in the case of 49 enzymes with more than 60 metabolism connections, the mean *K_e_* is 10.08±0.33, and the proportion of highly pleiotropic genes (*K_e_*≥9) is *F_P_*= 63% versus *F_P_*= 37% in the rest of enzymes (*P*<0.01). **Fig.4, B** shows a positive correlation between the number of biological processes and the mean of *K_e_* (*P*<0.001). We also observed that the fraction of highly pleiotropic genes with single GO (Gene Ontology) biological process is *F_P_*=0.22±0.008, which is mildly but significantly smaller than *F_P_*=0.25±0.011 among genes with multiple number of biological processes (*P*<0.05).

### The pleiotropy-connectivity correlation

Meanwhile, protein-protein interactions, enzyme connections, and involved biological processes together can only explain a small portion (∼6%) of gene pleiotropy variation among genes, raising some questions about the biological relevance of *K_e_*. Since current knowledge of gene multi-functionality is very limited, it would not be surprising that for most genes, the underlying molecular-cellular mechanism of gene pleiotropy remains unknown. Yet, for those highly pleiotropic genes, the percentage that can be biologically explained should be considerable. We thus examined this issue based on 803 highly pleiotropic genes with *K_e_*>9, and observed that about 21% highly pleiotropic genes can be explained by those hubs (high-connectivity) in the protein-protein interaction (PPI) network. By contrast, we used the low-pleiotropic gene group (674 genes for *K_e_*<6), and calculated that the percentage of hubs is ∼6.97% in the control group; randomization test has shown that the difference between the highly pleiotropic and lowly pleiotropic genes is highly significant (*P*<10^-6^). In short, we propose that the pleiotropy-connectivity is significant portion for highly pleiotropic genes, but spurious for lowly pleiotropic genes.

### Effect of baseline selection intensity

On the other hand, high expression level could increase the baseline selection intensity (*B*_0_) due to a variety of factors such as the translation efficiency, energy cost, or protein misfolding. For instance, the toxic effect of protein misfoldings after translation is proportional to the expression level. Hence, the constraint for sequence features that can reduce the rate of misfolding has nontrivial contributions to the baseline selection intensity but not to the gene pleiotropy. Indeed, we found *B*_0_ is correlated significantly with CAI or Abd (Table 1). However, it should be noticed that the expression effect seems nontrivial only for highly expressed genes, or genes with highly biased codon usages. In addition, the synonymous distance (*d_S_*) is highly correlated to *B*_0_ (*r^2^*=0.13, *P*<10^-10^), but virtually no correlation with the gene pleiotropy (*K_e_*) *(r^2^*=0.004). Hence, our current analysis reveals that both gene pleiotropy and the baseline selection intensity (*B*_0_) have nontribal contributions to the correlation between the expression level and the rate of protein evolution; yet the detail of the pattern remains further investigation.

### Toward a unified pleiotropy hypothesis

In summary, protein-protein interactions, expression broadness, enzyme connections, and involved biological processes are all molecular mechanisms for a gene to be pleiotropic, which explain well the positive correlations of gene pleiotropy with these genomic measures. After careful examinations, we are confident that our main results are robust under various technical treatments. Under this model, gene pleiotropy (*K*), which quantifies the multi-dimensionality of gene function in biological networks and systems, takes a pivotal role in protein evolution. However, current genomic data for gene multi-functionality together can only explain a small portion (<10%) of gene pleiotropy. We interpret this as the contribution of each genomic factor to gene pleiotropy is small and highly heterogeneous. Our conclusions differ from the expression hypothesis (Pal et al 2006; Koonin and Wolf 2006; Drummond et al. 2005; 2006; Subramanian and Kumar 2004; Salathe et al. 2006; Wolf et al. 2006) that gene expression is the primary factor for the rate of protein evolution. Alternatively, the pleiotropy hypothesis suggests that the rate variation among proteins is mainly due to the variation of gene pleiotropy, revealing a sophisticated display of gene functionality. Meanwhile, the baseline of protein sequence evolution may be determined by the biophysical-structure properties of proteins (DePristo et al. 2005; Koehl and Levitt 2002). Our study highlights the point that the evolutionary systems biology (Gerhart and Kirschner 1997; Pal et al 2006; Koonin and Wolf 2006) has becoming an important approach to understanding important biological problems.

## Data and Methods

### Human and mouse RNA-seq data

Human and mouse RNA-seq data were obtained from ###. We used the medium expression value among biological replicates to measure the expression level of a gene, following the common procedure of normalization. In this study, we use TPM>1 as the threshold for calling a gene “expressed in a given tissue”. The expression breadth of each gene was represented by the number of tissues this gene expressed. Using more stringent criteria would give a lower estimate of expression breadth, but our main results were essentially the same. Brain-preference genes are defined as those genes for which the highest expression is found in one of the brain tissues and this highest expression is at least twice the expression level in any non-brain tissue we have examined. Other tissue-tissue genes were identified similarly.

### Nuclear-encoding mitochondrial proteins

A collection of more than >1000 human nuclear-encoding mitochondrial genes were obtained from (Calvo, *et al*. (2006), which were predicted by the software Maestro with a 10% estimated false prediction rate. Finally 468 mitochondrial genes can be mapped to our data, and among of them, 19 mitochondrial ribosomal proteins were further curated.

### Protein interaction data sets

The high-throughput protein-protein interaction dataset (HTP-PI) used in the main context was created by union of four HTP studies (Utez et al. 2000; Gavin et al. 2002; Ho et al. 2002; Ito et al. 2001). 2677 of 4478 proteins in the HTP-PI dataset were mapped to yeast pleiotropic data. The LC-PI dataset (literature-curated protein interaction) (Reguly et al. 2006) was obtained from the BioGRID database (http://www.thebiogrid.org). 2122 of 3289 proteins in the LC-PI dataset mapped to yeast pleiotropic data. Another HTP dataset (Krogan’s data) was obtained from (Krogan et al. 2006), which was identified by tandem affinity purification (TAP) of affinity-tagged proteins expressed from their natural chromosomal locations followed by mass spectrometry.

### Analytical pipeline for estimating effective gene pleiotropy

*GenPleio.* We developed a computational pipeline (GenPleio) to estimate the effective number of molecular phenotypes (*K_e_*) and the effective selection intensity (*S_e_*). The underlying evolutionary framework and detailed procedure were described in our previous work (Gu 2007; Su et al 2010; Zeng et al. 2010; Gu 2014). Briefly, the estimation procedure was as follows. (*i*). Infer the phylogenetic tree from a multiple alignment of homologous protein sequences. Although there is no methodological preference, we require that the inferred tree topology should be roughly the same over methods. (*ii*). Estimate the nonsynonymous to synonymous ratio (*d_N_*/*d_S_*) from closely related coding sequences that satisfy *d_N_*/*d_S_* <1. (*iii*). Estimate the *H*-measure for rate variation among sites: Use the method of Gu and Zhang (1997) to infer the (corrected) number of changes at each site, under the inferred phylogeny. (*iv*). Estimate the *K_e_* and *S_e_* based on the nonsynonymous to synonymous ratio and *H*. The software Genpleio is available under request.

